# Buprenorphine exposure alters the development and migration of interneurons in the cortex

**DOI:** 10.1101/2020.10.26.356022

**Authors:** Vanesa Nieto-Estévez, Jennifer J. Donegan, Courtney McMahon, Hannah B. Elam, Teresa A. Chavera, Parul Varma, Kelly A. Berg, Daniel J. Lodge, Jenny Hsieh

## Abstract

The misuse of opioids has reached epidemic proportions over the last decade, with over 2.1 million people in the U.S. suffering from substance use disorders related to prescription opioid pain relievers. This increase in opioid misuse affects all demographics of society, including women of child-bearing age, which has led to a rise in opioid use during pregnancy. Opioid use during pregnancy has been associated with increased risk of obstetric complications and adverse neonatal outcomes, including neonatal abstinence syndrome. Currently, opioid use disorder in pregnant women is treated with long-acting opioid agonists, including buprenorphine. Although buprenorphine reduces illicit opioid use during pregnancy and improves infant outcomes at birth, few long-term studies of the neurodevelopmental consequences have been conducted. The goal of the current experiments was to examine the effects of buprenorphine on the development of the cortex using fetal brain tissue, 3D brain cultures, and rodent models. First, we demonstrated that we can grow cortical and subpallial spheroids, which model the cellular diversity, connectivity, and activity of the developing human brain. Next, we show that cells in the developing human cortex express the nociceptin opioid (NOP) receptor and that buprenorphine can signal through this receptor in cortical spheroids. Using subpallial spheroids to grow inhibitory interneurons, we show that buprenorphine can alter interneuron development and migration into the cortex. Finally, using a rodent model of prenatal buprenorphine exposure, we demonstrate that alterations in interneuron distribution can persist into adulthood. Together, these results suggest that more research is needed into the long-lasting consequences of buprenorphine exposure on the developing human brain.

## INTRODUCTION

The Opioid Epidemic has reached epic proportions, with over 2.1 million people in the US suffering from opioid use disorder (OUD) (1). This epidemic affects all demographics of society, including women of childbearing age. In a 2017 study, 6.5 percent, or 21,000, pregnant women, reported illicit use of prescription opioids (2, 3), which has been associated with obstetric complications, such as preeclampsia and premature labor (4–6), and adverse neonatal outcomes, including neonatal abstinence syndrome, an array of symptoms caused by dysregulation of central, autonomic, and gastrointestinal systems (7–9). To prevent these negative outcomes, the American Academy of Pediatrics and American College of Obstetricians and Gynecologists (ACOG) recommends opioid maintenance therapy to treat opioid use disorder during pregnancy. Buprenorphine, a partial mu- and kappa-opioid receptor agonist, is used for outpatient opioid maintenance therapy, and is the first-line treatment for OUD in pregnant women recommended by ACOG (10). Buprenorphine has been shown to be effective in reducing illicit opioid use in pregnant women (11) and improves infant outcomes at birth compared to other opioid maintenance therapies, including methadone (12). However, buprenorphine can cross the placental barrier and directly affect the developing neonate (13). While some studies have found that *in utero* buprenorphine exposure does not affect early childhood growth, cognitive development, language abilities, or sensory processing (14), others indicate that buprenorphine may have adverse effects on the developing fetus, including increased risk of prematurity and congenital malformations (14). Further, behavioral consequences have been observed in preschool-aged children, including motor, memory, and attention deficits, along with hyperactivity and impulsivity (14, 15). These conflicting results suggest that more research into the neurodevelopmental consequences of *in utero* buprenorphine exposure are urgently needed.

Studying fetal brain development in humans is extremely difficult due to ethical, time, and cost limitations. Further, multiple confounding factors, including obstetric complications and postnatal maternal care can have a major impact on the results. Therefore, in the current experiments, we use human pluripotent stem cells to generate 3D brain spheroids, which retain more of the structural and functional properties of the developing human brain (16–18). To generate excitatory neurons of the cortex, we grew cortical spheroids (hCS), which have ventricle-like structures surrounded by neuronal layers characteristic of the developing human cortex (17). To generate inhibitory interneurons, we grew subpallial spheroids (hSS), which express markers of developing interneurons (18, 19). These hCS and hSS were fused to examine the effect of buprenorphine exposure on cortical network development. Further, a rodent model was also used to confirm our *in vitro* findings and determine the long-term consequences of prenatal buprenorphine exposure.

In the current studies, we demonstrate that we can generate hCS and hSS, which resemble the developing human cortex and subpallium. We then show that in hCS, buprenorphine signals through the nociceptin opioid peptide (NOP) receptor, which is also expressed throughout the human fetal cortex. We also found that buprenorphine can alter markers of interneuron development in the hSS. When we fused the two spheroid subtypes, buprenorphine altered interneuron migration into the hCS and increased network activity. In our rodent model of prenatal opioid exposure, we observed changes in interneuron distribution throughout the cortex after buprenorphine exposure, suggestive of long-lasting alterations in interneuron migration. Together, these results suggest that buprenorphine exposure can alter cortical development and produce persistent changes in the cortical network.

## MATERIALS AND METHODS

### Pluripotent Stem Cell Generation and Maintenance

Human induced pluripotent stem cells (hiPSCs) were generated from a healthy 35-year-old male subject using an episomal reprogramming kit (Invitrogen). H9 ESCs were purchased from Wicell. Both hiPSCs and ESCs showed pluripotency markers and normal karyotype throughout the study (Figure S1A-B) and were grown on matrigel-coated plates in mTeSR medium (Stem Cell Technology) containing the Rock inhibitor, Y27632 (10uM, Selleck). In order to fluorescently label cells (Figure S1C), the Neon Transfection Kit (Invitrogen) was used to transfect cells with CRISPR-mCherry (mCh) plasmids. Cells were dissociated using accutase, then 1 x 10^6 cells were resuspended in Buffer R containing 1μg of CRISPR plasmid (AAVS1-T2 CRISPR in pX330) and 3 μg mCh plasmid (AAVS1-Pur-CAG-mCherry). Transfections were performed at 1100 V, 20 msec, 2 pulses and cells were immediately re-plated in mTeSR medium containing Y27632. Three days after transfection, 0.3 μg/ml puromycin was added to the media for 5 days to kill non-transfected cells.

### Drugs

Buprenorphine (2 ng/ml) was purchased from Cayman Chemical Company and oxycodone (20 ng/ml) was purchased from Spectrum Chemicals.

### Spheroid Cell Culture

To grow hCS and hSS we used the method described by Pasca and colleagues with light modifications (Pasca et al 2015, Birey et al 2017). Briefly, 9,000 stem cells were resuspended in 150 μl neural induction medium (DMEM/F12, 20% knockout serum replacement, Glutamax, MEM-NEAA, 0.1mM 2-mercaptoethanol, Peni/Strep) containing Y27632 (20 μM). Media was changed daily. On days 1-5, the SMAD inhibitors, dorsomorphin (5 μM, Sigma) and SB-431542 (10 μM, Tocris) were added. On days 6-24, media was replaced with neural medium (Neurobasal A, B-27 without Vitamin A, Glutamax, Peni/Strep) containing the growth factors, bFGF (20 ng/ml, Peprotech) and EGF (20 ng/ml, Peprotech). On day 25, spheroids were transferred to neural medium containing BDNF (20 ng/ml, Peprotech) and NT-3 (20 ng/ml, Peprotech) until day 42. hSS also received the Wnt pathway inhibitor, IWP-2 (5 μM, Selleckchem) on days 4-22 and the Smo pathway activator, SAG (100 nM, Selleckchem) on days 12-22 (Figure 1A).

**Figure 1.**
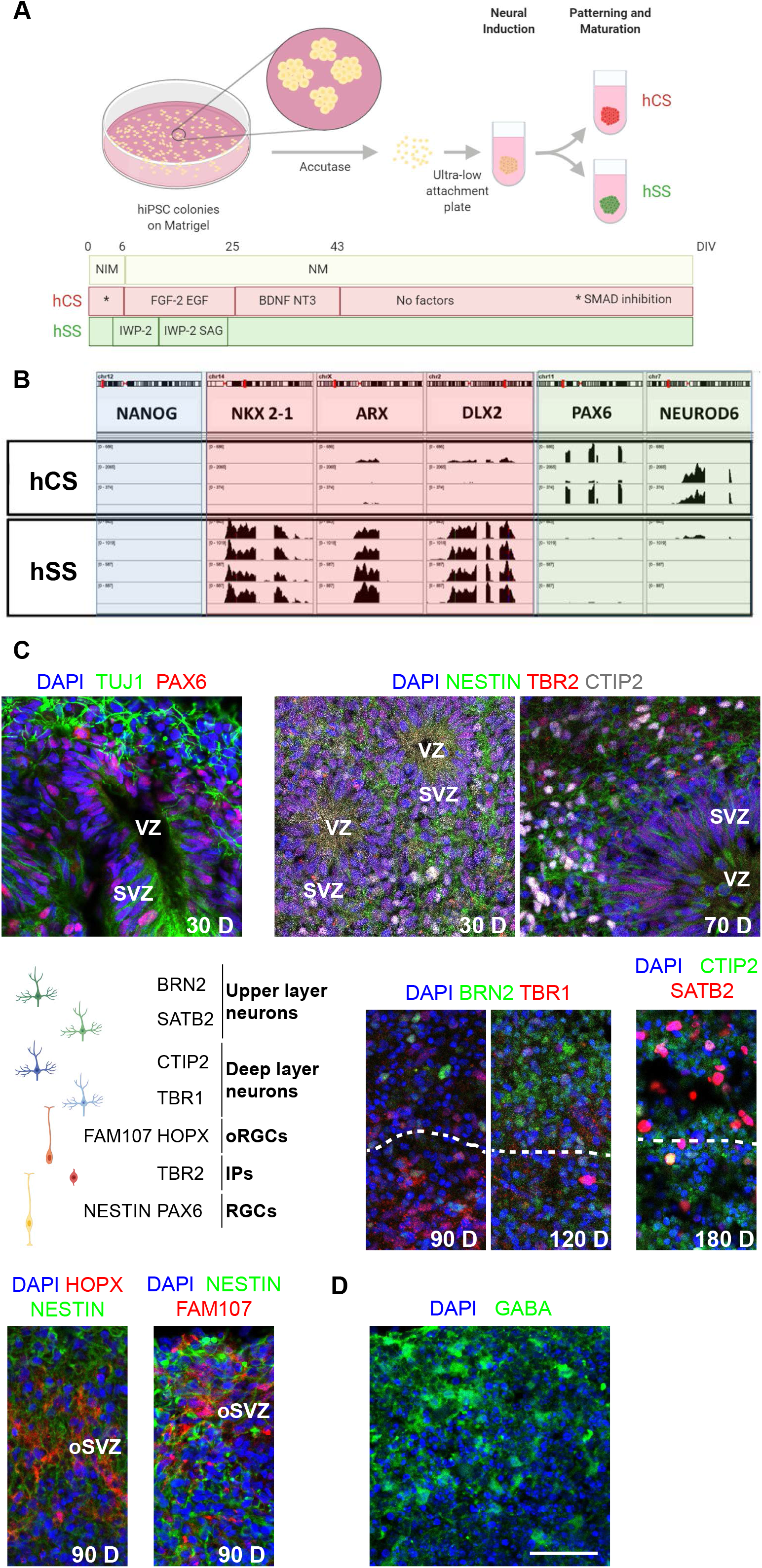
Characterization of cortical and subpallial spheroids. The protocol for generating cortical and subpallial spheroids is depicted in (A). B) RNA Sequencing found that neither cortical nor subpallial spheroids express *NANOG*. Cortical spheroids express *PAX6* and *NEUROD6* while subpallial spheroids express *NKX2-1, ARX*, and *DLX2*. n=3-4. C) Immunohistochemistry demonstrating that cortical spheroids express ventricular-like zones and express markers of cortical development. n=6 D) Subpallial spheroids express GABA. n=6 Scale bar = 50 μm

### RNA Sequencing

On day 60, hCS and hSS spheroids were homogenized and RNA was prepared using the Qiagen RNEasy Plus Micro Kit according to manufacturer’s instructions. Library construction, sequencing, and analysis were performed by NovoGene.

### Spheroid Immunohistochemistry

Spheroids were rinsed in DPBS, then fixed in 4% paraformaldehyde for 30 minutes - overnight at 4°C. After rinsing in DPBS, spheroids were incubated in 30% sucrose for 24-48 hours at 4°C. Spheroids were embedded in OCT compound and frozen. A cryostat was used to cut 14-25 micron sections, which were mounted on gelatin-coated slides. For immunohistochemistry, slides were washed in PBS containing 0.3% Triton X-100, then blocked with 3-5% normal goat serum for 1 hour. Slides were incubated in primary antibody (Table 1) overnight at 4°C in a humidified chamber. After washing, slides were incubated in secondary antibody (Jackson Immunoresearch, 1:500) for 1-2 hours at room temperature. Slides were washed, DAPI (Sigma) was added to label nucleus, then coverslipped using Poly-Vinyl Alcohol (Sigma). Sections were imaged using a Leica Microscope (TCS SPE8) and fluorescence intensity was analyzed using Image J software.

**Table 1.**
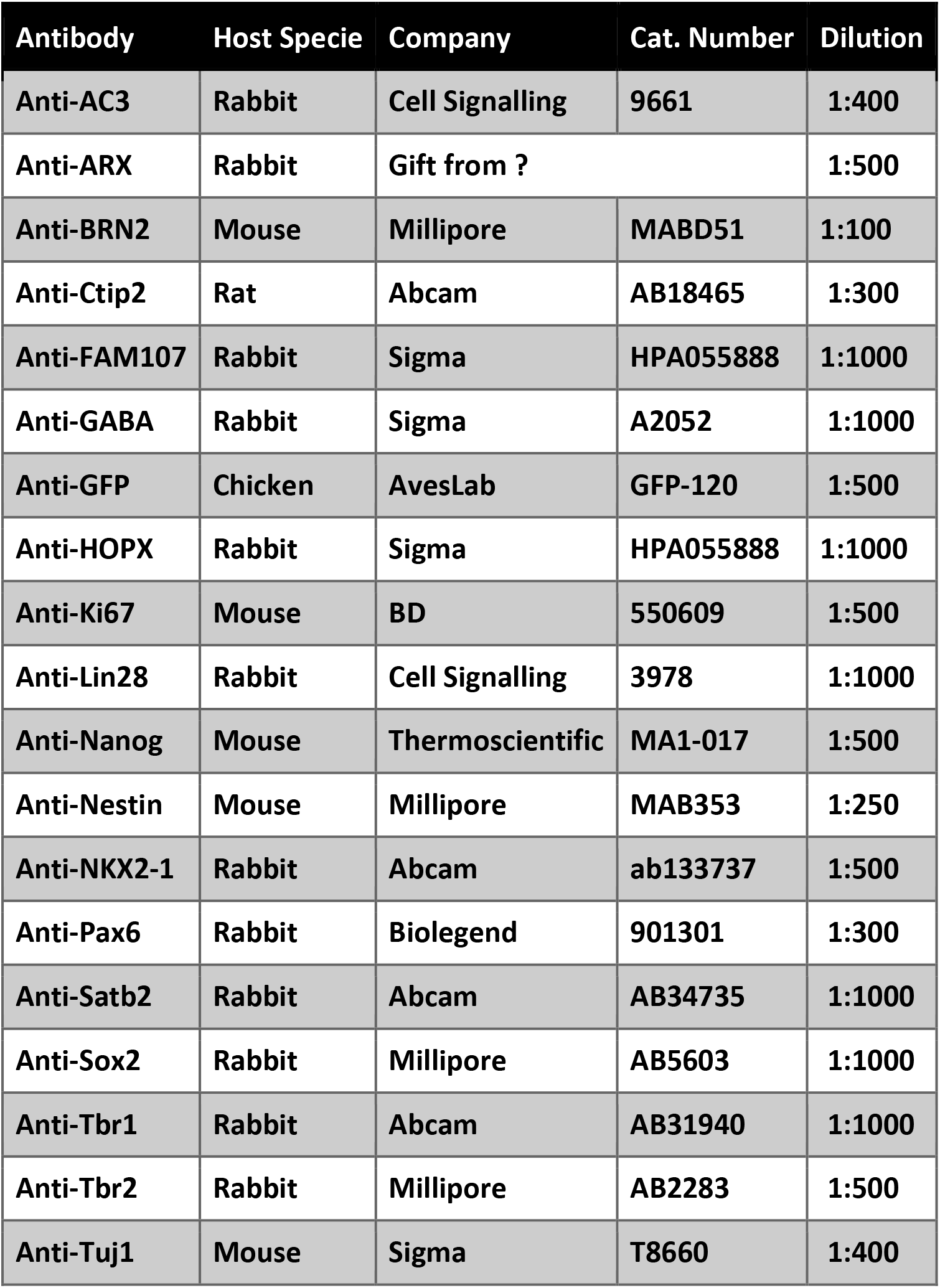

| Antibody | Host Specie | Company | Cat. Number | Dilution |
| --- | --- | --- | --- | --- |
| Anti-AC3 | Rabbit | Cell Signalling | 9661 | 1:400 |
| Anti-ARX | Rabbit | Gift from ? |  | 1:500 |
| Anti-BRN2 | Mouse | Millipore | MABD51 | 1:100 |
| Anti-Ctip2 | Rat | Abcam | AB18465 | 1:300 |
| Anti-FAM107 | Rabbit | Sigma | HPA055888 | 1:1000 |
| Anti-GABA | Rabbit | Sigma | A2052 | 1:1000 |
| Anti-GFP | Chicken | AvesLab | GFP-120 | 1:500 |
| Anti-HOPX | Rabbit | Sigma | HPA055888 | 1:1000 |
| Anti-Ki67 | Mouse | BD | 550609 | 1:500 |
| Anti-Lin28 | Rabbit | Cell Signalling | 3978 | 1:1000 |
| Anti-Nanog | Mouse | Thermoscientific | MA1-017 | 1:500 |
| Anti-Nestin | Mouse | Millipore | MAB353 | 1:250 |
| Anti-NKX2-1 | Rabbit | Abcam | ab133737 | 1:500 |
| Anti-Pax6 | Rabbit | Biolegend | 901301 | 1:300 |
| Anti-Satb2 | Rabbit | Abcam | AB34735 | 1:1000 |
| Anti-Sox2 | Rabbit | Millipore | AB5603 | 1:1000 |
| Anti-Tbr1 | Rabbit | Abcam | AB31940 | 1:1000 |
| Anti-Tbr2 | Rabbit | Millipore | AB2283 | 1:500 |
| Anti-Tuj1 | Mouse | Sigma | T8660 | 1:400 |

### qPCR

hCS and hSS were treated with vehicle, buprenorphine or oxycodone for 10 days. On day 60, spheroids were homogenized and RNA was prepared using the Qiagen RNEasy Plus Micro Kit according to manufacturer instructions. RNA concentration and quality were determined using a NanoDrop, then total RNA was converted to cDNA using the Applied Biosystems High Capacity Reverse Transcription Kit. Real-time quantification of diluted cDNA was performed in triplicate reactions containing sample (4 ng), IDT Primetime Gene Expression Master Mix (2X), and IDT Primetime qPCR Assay (20X) on a BioRad CFX384 Real Time System. Cycling Conditions consisted of one cycle at 95°C for 3 min, followed by 40 cycles of denaturation (95°C for 15 sec) and elongation (60°C for 1 min). The relative gene expression was calculated using the 2^-ΔΔCT^ method. The following IDT Primetime Gene Expression Assays were used: Dlx1 (Hs.PT.58.4632198), Dlx5 (Hs.PT.58.45646524), Lhx6 (Hs.PT.58.27682011), Arx (Hs.PT.58.213.7622), Nkx2.1 (Hs.PT.58.2461055), ADCY3 (Hs.PT.569.2298693), BCL11B (Hs.PT.58.27217530), MKi67 (Hs.PT.58.27920212), Pax6 (Hs.PT.58.25914558), SATB2 (Hs. PT. 58.24560574), TBR1 (Hs.PT.58.24815929), TUBB3 (Hs.PT.58.20385221) and GAPDH (Hs.PT.39a.22214836).

### cAMP Assay

NOP receptor-mediated inhibition of adenylyl cyclase activity was determined by measuring cAMP accumulation in the presence of the adenylyl cyclase activator, forskolin, and the phosphodiesterase inhibitor, rolipram as previously described (20). Briefly, spheroids were incubated in wash buffer (HBSS containing 20 mM HEPES, pH 7.55) for 15 min at 37°C. Rolipram (100 μM, Company) was added along with the following drugs: buprenorphine (5 nM), nociception (150 nM, Tocris), or oxycodone (60 nM) in the presence or absence of the NOP Receptor antagonist, UFP-101 (10 μM, Tocris). Forskolin (10 μM, Company) was then added and spheroids were incubated for an additional 15 min. Incubation was terminated by aspiration of the wash buffer and addition of 500 μl of ice-cold ethanol. The ethanol extracts from individual spheroids were dried under a gentle airstream and reconstituted in 100 μl of sodium acetate (pH 6.2) and the cAMP content was determined by radioimmunoassay (RIA).

### Multi-Electrode Array (MEA) Assay

To record network activity, the Harvard MCS Multiwell-MEA-system, which contains a 12-electrode grid in each well of a 24 well plate, was used. Twenty-four hours before the MEA was conducted, spheroids were transferred to the MEA plate containing BrainPhys media (STEMCELL Technologies). For acute experiments, a 10-minute baseline recording was followed by treatment with buprenorphine, or oxycodone in the presence of vehicle or the NOP receptor antagonist, UFP-101. Fifteen minutes later, 3 10-minute recordings were performed. For the chronic experiments, hCS or fused hCS-hSS spheriods were treated with vehicle, buprenorphine, or oxycodone for 10 days. On day 60 or 120, spheroids were transferred to MEA plates containing BrainPhys media (drugs were not present) and 3 10-minute recordings were performed. Data was sampled at 20 kHz, digitized and analyzed using the Harvard Multiwell-Analyzer software with a 100 Hz high pass and 3.5 kHz low pass filter and an adaptive spike detection threshold was set at 8 times the standard deviation for each electrode. The results from each 10-minute recording were averaged.

### Interneuron Migration

On day 30, hSS were infected with Lenti hDlx1/2b:GFP (gift from Dr. John Rubenstein). On day 40, one hSS and one hCS were transferred to an Eppendorf tube containing neural media and incubated without disruption for 3-4 days. After fusion was complete, fused spheroids were treated with buprenorphine or oxycodone from days 50-60. On day 60, spheroids were transferred to glass-bottom (Corning) plates and the migration of hDlx1/2b:GFP interneurons into cortical tissue was recorded. Time-lapse images were taken every 20 min for 24 hours using a Leica Microscope (TCS SPE8) (Figure 4A). Analysis of interneuron migration was performed using Imaris imaging software.

### Rat Experiments

Pregnant female rats were injected with 1 mg/kg buprenorphine or 10 mg/kg oxycodone on gestational days 11-21. Male and female pups were weaned on postnatal day 21 and all experiments were performed in adult animals (after postnatal day 42; Figure 5A). Animals were maintained in a temperature-controlled environment on a 14:10 h light-dark cycle and had access to food and water *ad libitum*. All procedures were consistent with NIH guidelines (NIH publications no. 80-23, revised 1978) and approved by the Institutional Animal Care and Use Committee of the University of Texas Health Science Center at San Antonio.

### Rat Immunohistochemistry

Adult rats were transcardially perfused with saline followed by 4% paraformaldehyde. Brains were post-fixed and cryopreserved for at least 24 hours. Coronal sections (50 μm) through the medial prefrontal cortex were blocked (5% normal goat serum), incubated with a mouse anti-Lhx6 primary antibody (Santa Cruz, 1:250), then washed and incubated with a goat anti-mouse Alexafluor488 secondary antibody (Invitrogen). Sections were mounted, coverslipped with Prolong Gold Antifade Reagent (Molecular Probes), then imaged using a Zeiss AxioObserver inverted microscope. To analyze the migration of interneurons into the cortex, 10 equidistant bins through the medial prefrontal cortex were defined to determine the distribution of cells across the cortical layers (Meechan PNAS). Lhx6+ cells were counted in at least 3 sections per animal by a blind experimenter.

### Human Fetal Brain Tissue Analysis

Fetal brain bulk RNA sequencing dataset was downloaded from BrainSpan: Atlas of the Developing Human Brain (https://brainspan.org/). RPKM (reads per kilobase transcripts per million mapped reads) values for OPRL1 gene were mined and normalized for each sample using the following formula RPKM(gene)/(∑RPKM(all genes in the sample)) X 10^6. Graphpad Prism was used to plot OPRL1 normalized counts from 8-37 weeks post conception and median values were used to draw conclusions about the temporal expression pattern.

### Statistical Analysis

In all figures, data are shown as mean + s.e.m and n is indicated in the figure legend. Data was analyzed by one- or two-way ANOVA and the Holm-Sidak post-hoc test was used when significant interactions were present. When comparing groups with unequal variances the nonparametric Kruskal-Wallis test was used followed by Dunn’s multiple comparison test. All tests were two-tailed, and significance was determined at p<0.05.

## RESULTS

### Generation of cerebral and subpallial spheroids

To study the effect of buprenorphine during human brain development, we first generated cortical spheroids (hCS) and subpallial spheroids (hSS) from hiPSCs and ESCs using the method described by Pasca and colleagues (Fig 1A) (17, 18). Both cell lines expressed pluripotent stem cell markers and maintained normal karyotype (Supp Fig 1). RNA sequencing was used to confirm spheroid identity. Neither cortical spheroids nor subpallial spheroids expressed *NANOG*, a homeobox protein involved in maintaining embryonic stem cell pluripotency (Fig 1B) (21). Cortical spheroids expressed *PAX6*, a marker for dorsal forebrain progenitor cells (22), and *NeuroD6*, a gene expressed ubiquitously throughout the fetal cerebral cortex (Figure 1B) (23). These genes were not expressed in subpallial spheroids. Conversely, subpallial spheroids expressed *ARX*, *DLX2*, and *NKX2-1* (Fig 1B). Dlx2 has been shown to bind to *ARX* and regulate the migration of GABAergic neurons (24). Similarly, *NKX2-1* is a homeobox transcription factor that regulates the fate specification and migration of GABAergic interneurons (25). These subpallial markers were not observed in hCSs. In addition, hCS showed VZ-like structures at early stages, including radial glia and intermediate progenitor cells expressing PAX6, NESTIN, and TBR2. At later time points, we observed neurons of both deep (TBR1+ and CTIP2+) and upper layers (SATB2+ and BRN2+). We also found markers of outer radial glial cells such as HOPX and FAM107 (Fig 1C). Moreover, hSS presented GABA+ cells (Fig 1D). These data showed hCS and hSS resembled features of cortical and subpallial brain development *in vitro*.

### Buprenorphine signals through the nociception opioid peptide receptor in cortical spheroids

Buprenorphine is thought to have its therapeutic effect via the mu- and kappa-opioid receptors; However, buprenorphine also acts as a full agonist at the nociceptin opioid peptide (NOP) receptor (26). Therefore, expression of these opioid receptors was determined in hCS and hSS using RNA sequencing. In line with the pattern of opioid receptor expression in the human fetal brain, neither the mu-, kappa-, nor delta-opioid receptors were significantly expressed in either the cortical or subpallial spheroids (Fig 2A). However, the nociceptin opioid peptide (NOP) receptor is expressed in both cortical and subpallial spheroids (Fig 2A). In addition, we confirmed the expression of NOP receptor in human fetal tissue using publicly available dataset downloaded from BrainSpan, Atlas of the Developing Human Brain (https://brainspan.org/) (Fig 2B). After confirming NOP receptor expression, cAMP assays and multielectrode arrays were performed on Day 50 cortical spheroids (Fig 2C) to determine if buprenorphine can have functional effects on the spheroids via the NOP receptor. Activation of the NOP receptor, which typically signals through Gi proteins, leads to an inhibition of adenylyl cyclase activity and decreases in cellular cAMP levels. Therefore, cAMP assays were first used to determine whether buprenorphine can activate intracellular signaling pathways via the NOP receptor. We found that buprenorphine decreased cAMP accumulation compared to vehicle-treated spheroids (Fig 2D, Two-way ANOVA: Interaction F(3,68)=3.33, p<0.05; Opioid F(3,68)=11.90, p<0.05; NOP Antagonist F(1,68)=50.75, p<0.05; Holm-Sidak Post-Hoc Test: Vehicle-Vehicle vs Buprenorphine-Vehicle t=3.42, p<0.05). These results are in line with the ability of nociceptin, the endogenous NOP receptor ligand, to decrease cAMP accumulation (Fig 2C, Holm-Sidak Post-Hoc Test: Vehicle-Vehicle vs Nociceptin-Vehicle t=5.48, p<0.0-5). Pretreatment with the NOP receptor antagonist UFP-101 completely reversed buprenorphine- and nociceptin-induced decrease in cAMP accumulation (Fig 2D, Buprenorphine-Vehicle vs Buprenorphine-UFP t=3.16, p<0.05, Nociceptin-Veh vs Nociceptin UFP t=6.32, p<0.05), confirming that in hCS, buprenorphine can signal through the NOP receptor.

**Figure 2.**
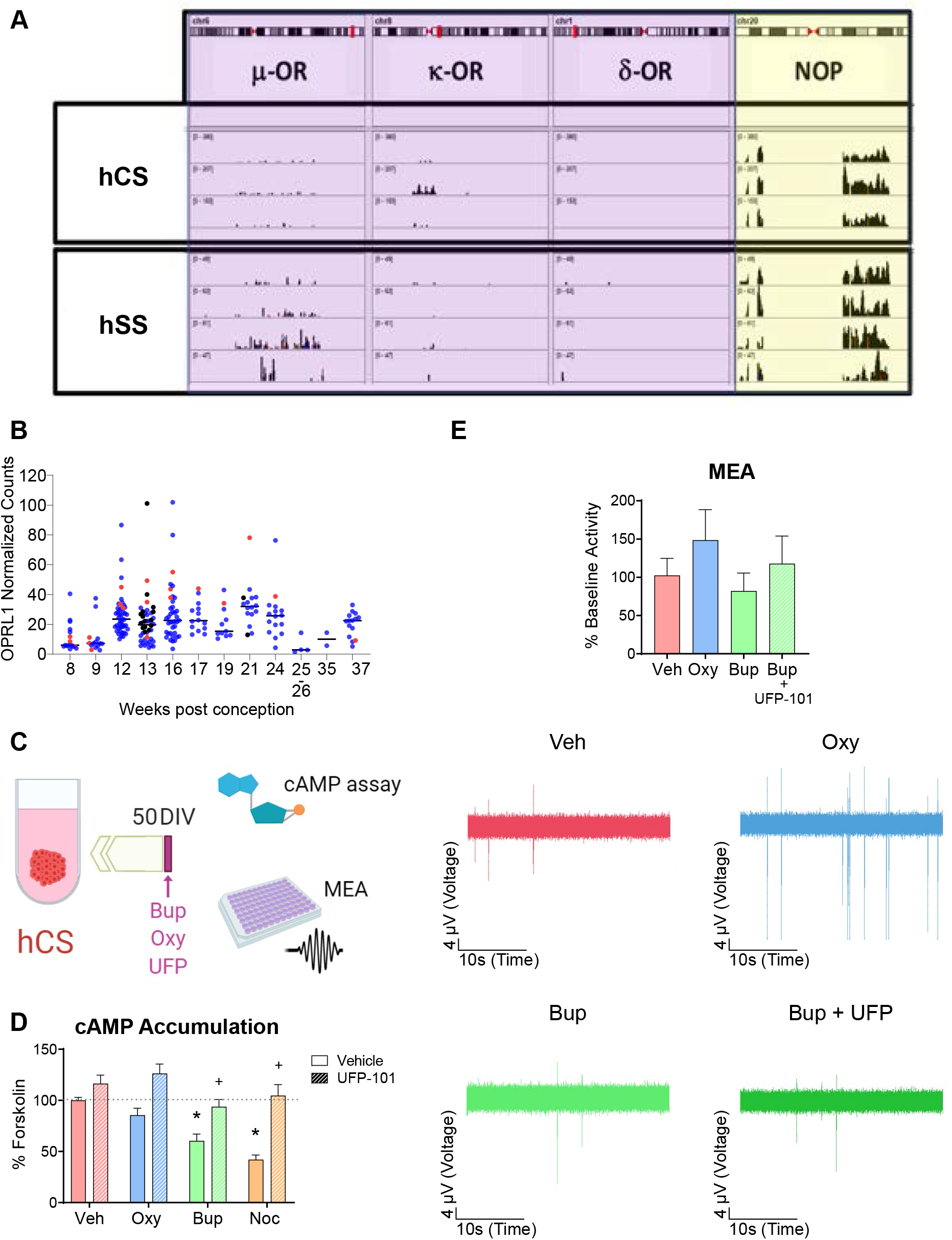
Buprenorphine signals through the nociception receptor. A) RNA Sequencing found that both cerebral and subpallial spheroids express the NOP receptor but not the mu-, kappa- or delta-opioid receptors. n=3-4. B) Data from fetal brain tissue demonstrating NOP receptor expression in cortical and subpallial regions beginning at 12 weeks. Red dots indicate ganglionic eminences and striatum, blue dots indicate cortical tissue. n=1-2. The experimental design is depicted in (C). D) Buprenorphine, like the endogenous NOP receptor ligand, decreases cAMP accumulation. This effect is blocked by treatment with the NOP receptor antagonist, UFP-101. n=8-12. E) Buprenorphine produces a trend toward decreased activity in the MEA, an effect that was blocked by UFP-101. n=16-21.

To determine whether buprenorphine’s inhibition of cAMP signaling was accompanied by a change in neural activity, MEA recording of cortical spheroids were then conducted to investigate changes in extracellular activity following acute exposure to the opioids. Compared to vehicle-treated spheroids, spheroids treated with buprenorphine acutely displayed a trend toward decreased activity following a 15-min exposure. Spheroids co-treated with UFP-101 and buprenorphine showed activity levels similar to that of vehicle-treated, further suggesting that buprenorphine acts upon the NOP receptor (Fig 2E, One-way ANOVA F(3,71)=0.85, p>0.05). To then determine if chronic opioid exposure affects cortical development, hCS were treated with buprenorphine or oxycodone for 10 days (day 50-60) and qPCR, immunohistochemistry, and MEAs were performed (Fig. S2 A). While chronic treatment with either buprenorphine and oxycodone altered expression of some genes in hCS (Fig. S2 B-D), neither drug produced significant changes in cortical activity (Fig S2 E).

### Buprenorphine alters markers of interneuron development in subpallial spheroids

After demonstrating that buprenorphine can influence the function of cortical spheroids via the NOP receptor, the effect of buprenorphine on subpallial spheroids was also determined. Day 60 subpallial spheroids were analyzed by qPCR and immunohistochemistry after a 10 day treatment with oxycodone or buprenorphine (Fig 3A). First, qPCR was used to measure mRNA expression of *ARX, DLX1, LHX6*, and *NKX2-1*, genes involved in the development and migration of interneurons (27). There was a main effect of treatment (Two-way ANOVA: Opioid F(2,164)=3.06, p<0.05), driven by the increase in interneuron gene expression caused by buprenorphine exposure. Then, we performed immunostainings against ARX, KI67 and NKX2.1. We found a significant effect of ARX+ cells in hSS after opioid exposure (Fig 3C-F). This data suggests that opioids can affect interneuron progenitors during brain development.

**Figure 3.**
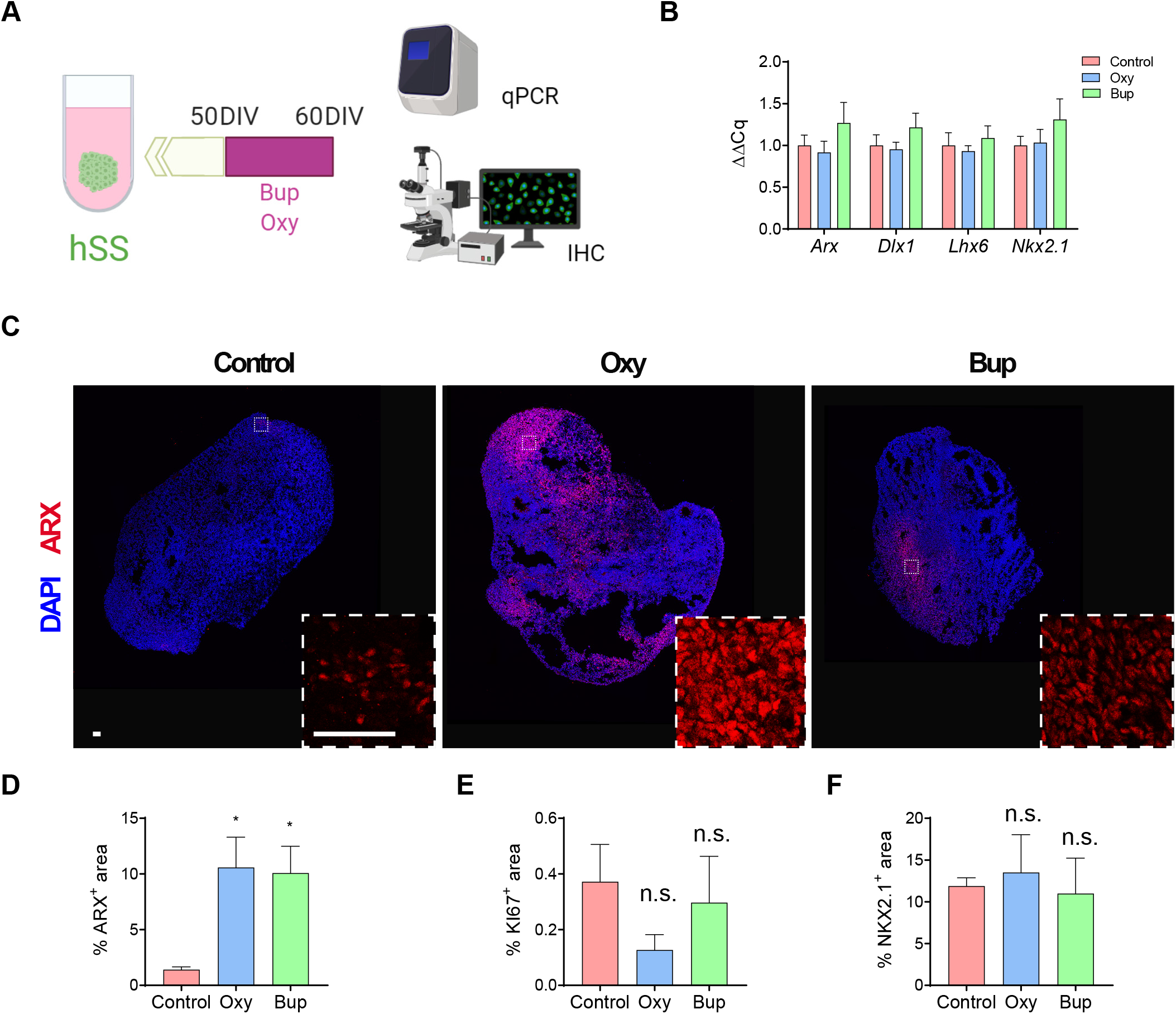
Buprenorphine alters markers of interneuron development in subpallial spheroids. The experimental design is depicted in (A). B) Buprenorphine treatment increases the expression of interneuron markers in subpallial spheroids. n=12-17. C) Both oxycodone and buprenorphine treatment increase ARX area in subpallial spheroids. KI67 and NKX2-1 were not affected. n=5-6. Scale bar = 50 μm.

### Buprenorphine alters interneuron migration and disrupts network activity in fused spheroids

Migration of interneurons from the ganglionic eminences into the cortex is essential to cortical development (27). In order to determine the effect of opioids on migrating interneurons, subpallial and cortical spheroids were fused and the migration of GFP-labeled interneurons were imaged over a 24 hour period (Fig 4A). There was a significant effect of opioid treatment on the percent of GFP-positive cells that migrated into the cortical spheroids from the subpallial area (Fig 4B-C; One-way ANOVA F=10.63, p<0.05). Post-hoc analysis demonstrated that buprenorphine produced a significant increase in the percentage of GFP-positive cells that migrated into the cortex compared to both control- and oxycodone-treated spheroids (Holm-Sidak: Control v Buprenorphine t=4.201; Buprenorphine v Oxycodone t=3.748). Next, we determined if this change in interneuron migration had an effect on network activity using MEA. Cortical spheroids were treated with vehicle, buprenorphine, or oxycodone for 10 days. At day 60, MEAs were conducted to determine the effect of chronic opioid exposure on the extracellular activity of the spheroids. Although not significant, Buprenorphine-treated spheroids exhibited a trend toward increased activity compared to the vehicle- and oxycodone-treated spheroids. (Fig 4D, One-way ANOVA F(2,11) = 0.89, p>0.05).

**Figure 4.**
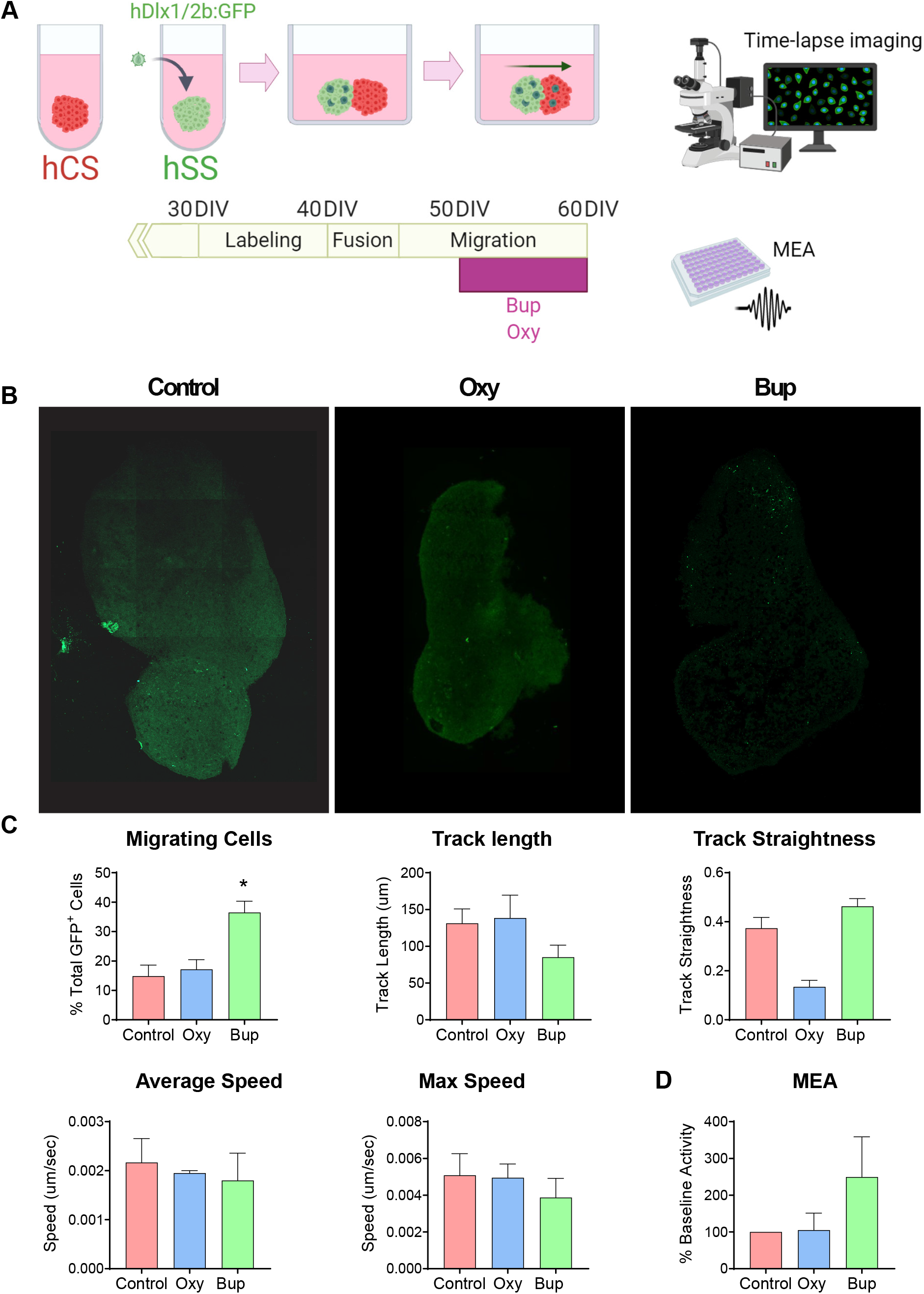
Buprenorphine alters interneuron migration into cortical spheroids and disrupts network activity. The experimental design is depicted in (A) Representative images of fused spheroids are shown in (B) C) Buprenorphine increased the number of GFP+ cells that migrated into the cortical spheroid compared to control- and oxycodone-treated spheroids. n=6. D) Buprenorphine exposure produces a trend toward increased MEA activity compared to control- and oxycodone-treated spheroids. n=6-7. Scale bar = TBD.

### Prenatal buprenorphine exposure alters interneuron migration in the rodent cortex

To determine if buprenorphine can also disrupt interneuron migration *in vivo*, pregnant rats were injected with buprenorphine or oxycodone on gestational days 11-21 (Fig 5A). In adult offspring, the total number and distribution of Lhx6+ interneurons were analyzed throughout the medial prefrontal cortex (mPFC; Fig 5B). Prenatal opioid exposure had no effect on the total number of Lhx6+ positive cells in the mPFC (Figure 5C; One-way ANOVA F=0.65, p>0.05). However, the laminar distribution of Lhx6+ cells was altered by prenatal buprenorphine exposure (Fig 5D-E; Mixed effects ANOVA Layer F(2.9,20.4)=67.45, p<0.05). Specifically, in bins 7 and 8, there were significantly more Lhx6+ cells in the buprenorphine-treated animals (Holm-Sidak Bin 7 t=-2.4, p<0.05; Bin 8 t=-3.9, p<0.05). Together, these results suggest that prenatal buprenorphine exposure can alter interneuron migration into the developing cortex, and the altered distribution of interneurons can persist into adulthood.

**Figure 5.**
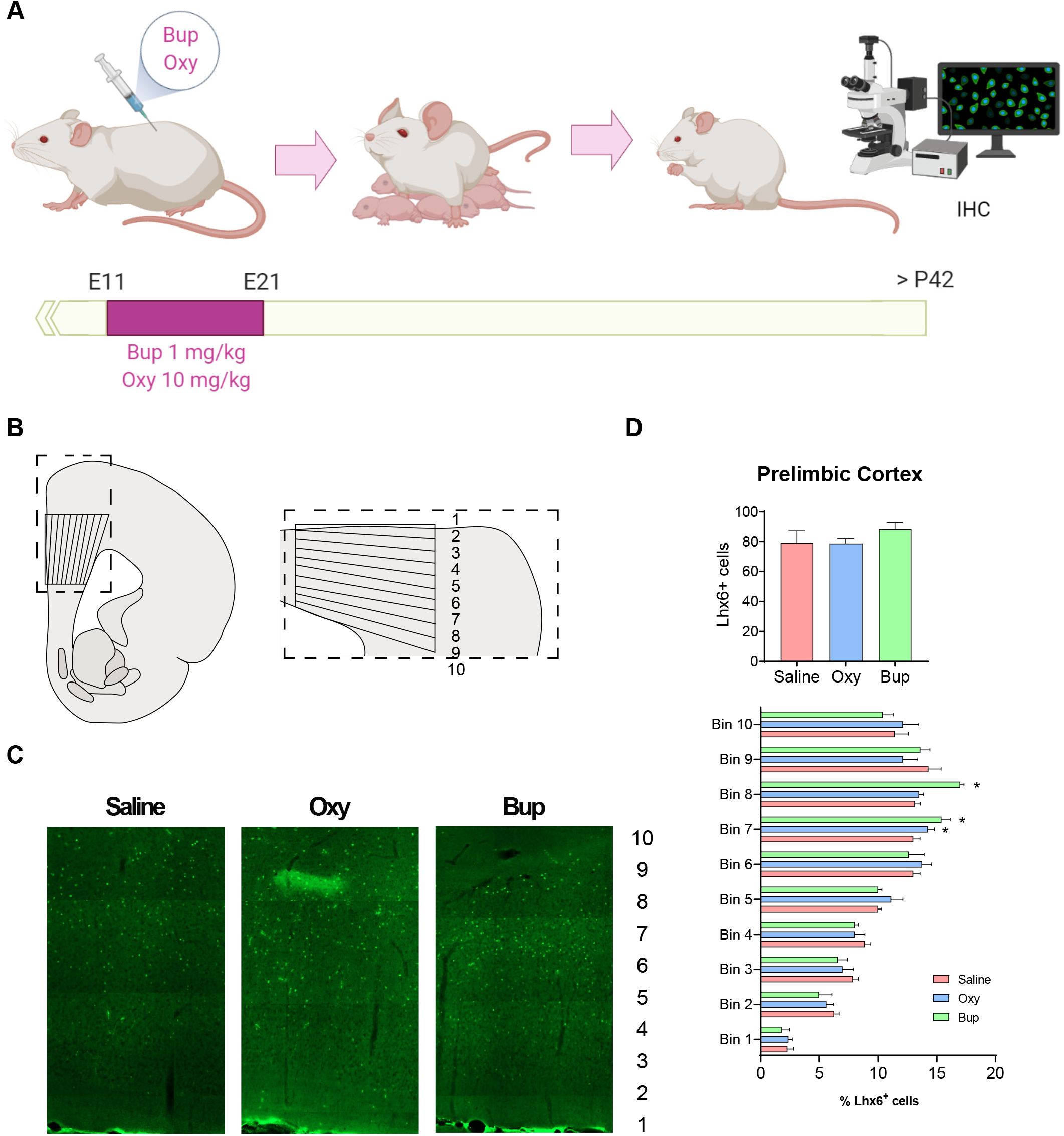
Prenatal buprenorphine alters interneuron migration in the rodent cortex. The experimental design is depicted in (A). The region of the medial prefrontal cortex analyzed is shown in (B). Representative images of Lhx6+ cells are shown in (C). D) The total number of Lhx6+ cells in the prelimbic region (PrL) of the mPFC is not altered by *in utero* opioid exposure. The distribution of Lhx6+ cells throughout the PrL is altered by buprenorphine exposure. n=5-8.

## DISCUSSION

In the current experiments, we used data from fetal brain tissue, human brain spheroids, and a rodent model to demonstrate that buprenorphine exposure during development can alter interneuron migration and disrupt cortical network activity via the NOP receptor. Buprenorphine, the preferred treatment for opioid use disorder (OUD) in pregnant women (10), acts as a partial agonist at mu- and kappa-opioid receptors to alleviate withdrawal symptoms while blocking the effect of exogenously administered opioids (26). However, this drug is also a full agonist at the nociceptin opioid peptide (NOP) receptor (26). The NOP receptor, encoded by the *OPRL1* gene, is a member of the opioid family of G protein-coupled receptors but is not activated by classical opioids with known abuse liability (28). Upon ligand binding, the NOP receptor acts through Galphai/Galphao to inhibit adenylate cyclase, activate mitogen-activated protein kinases, increase K+ conductance, and inhibit Ca2+ conductance (29, 30). Previous work has demonstrated that the NOP receptor is expressed throughout the developing rat and human brain, including in the neocortex (31). We confirmed these findings using a published RNA sequencing data set from human fetal brain tissue. Specifically, we found that the NOP receptor is expressed in cortical and subpallial fetal brain tissue as early as 12 weeks post-conception and remains elevated until birth, suggesting that buprenorphine treatment *in utero* has the potential to affect cortical development through its activity at the NOP receptor.

After demonstrating that the NOP receptor is present in fetal brain tissue, we then moved to an *in vitro* model to confirm that buprenorphine can signal through this receptor in developing cortical tissue. Specifically, we used human pluripotent stem cells to grow cortical spheroids, which resemble the developing cortex with ventricular zone-like structures, progenitor cells expressing PAX6, NESTIN, TBR2, and immature neurons that express TUJ1. Cortical spheroids also express cortical layer markers, including TBR1 and CTIP2 (deep layer neurons) and SATB2 and BRN2 (upper layer neurons). We used RNA sequencing to demonstrate that the NOP receptor is present in cortical spheroids. Interestingly, we did not observe expression of the mu-, k-, or d-opioid receptors in the human cortex, which is in line with results from fetal brain suggesting that these receptors are not expressed until later in development. Then, we used cAMP and MEA assays in these cortical spheroids to demonstrate that the NOP receptor is not only present, but that buprenorphine can signal via this receptor in developing brain tissue.

The cortex is made up of a variety of cell types, including interneurons, which are derived from a group of subpallial brain regions, the ganglionic eminences, before migrating tangentially into the cortex (27). Therefore, we also used human pluripotent stem cells to grow subpallial spheroids, which contain GABA-expressing neurons and express genes associated with interneuron development (*NKX2-1, ARX*, and *DLX2*). We found that chronic buprenorphine exposure increased markers of interneuron development in subpallial spheroids. Further, when cortical and subcortical spheroids were fused (19), we found that buprenorphine exposure altered interneuron migration, leading to an increase in interneuron migration into cortical spheroids. This increase in interneuron migration was accompanied by an increase in network activity in fused spheroids exposed to buprenorphine chronically. These results may seem contradictory as interneurons signal via the inhibitory neurotransmitter, GABA. However, high intracellular chloride concentrations in the developing brain can result in depolarization of the cell when GABA binds to its ionotropic receptors (32, 33). Interestingly, the excitatory action of GABA in the developing brain has been shown to induce synaptogenesis (34) and produce long-term changes in synaptic efficacy (35, 36), suggesting that alterations in the development of GABAergic interneurons by buprenorphine may shape the development of the cortex.

Network activity in the adult cortex is regulated by a complex interplay between excitatory and inhibitory neurons. Excitatory pyramidal cells release glutamate and are primarily responsible for transmitting information between brain regions. Conversely, inhibitory interneurons use GABA to signal and act locally to maintain control over individual pyramidal cells and to regulate oscillations across neuronal assemblies (37). After demonstrating that buprenorphine can alter interneuron migration in the developing cortex, we next used a rodent model of prenatal opioid exposure to show that buprenorphine exposure *in utero* produces changes in interneuron distribution throughout the adult cortex. These findings may be significant as interneuron dysfunction has been associated with a variety of neurodevelopmental and psychiatric disorders, including schizophrenia, autism, and intellectual disability (37). For example, schizophrenia patients have decreased expression of the GABA synthesizing enzyme, glutamic acid decarboxylase (GAD), in the prefrontal cortex, an effect that seems to reflect not a loss of cells but rather a loss of function in specific subclasses of interneurons (38). We have recently demonstrated that disrupting interneuron function in the cortex can produce schizophrenia-like deficits in social interaction and cognitive function in otherwise healthy animals (39). Further, our lab and others have shown that restoring interneuron function in the prefrontal cortex in a rodent developmental disruption model can attenuate behavioral deficits (40). Together with the current findings, these results suggest buprenorphine alters the development of cortical circuits implicated in the pathology of neurodevelopmental and psychiatric disorders.

Interestingly, the NOP receptor and its endogenous ligand, nociceptin, have also been implicated in psychiatric disease (41, 42). In adults, the receptor is highly expressed in brain regions associated with mental illness, including the cortex, hippocampus, amygdala, thalamus, hypothalamus, and dorsal raphe (31, 43–46). Further, nociceptin levels are elevated in patients suffering from bipolar disorder and major depression (47). A recent proof of concept clinical trial demonstrated that the NOP receptor antagonist, LY2940094, improved depression scores compared to placebo (41). These limited human findings are corroborated by animal models. Specifically, antidepressant-like effects in the forced swim test have been observed in both NOP receptor knock-out animals (48) and in wild-type rodents treated with a NOP receptor antagonist (41, 49). Activation of the NOP receptor has also been shown to impair working memory, one cognitive function associated with schizophrenia (50). While the current experiments did not examine how buprenorphine exposure *in utero* may affect NOP receptor function in adulthood, previous work has demonstrated that in rodents, prenatal buprenorphine exposure can decrease G protein coupling to NOP receptors (51). Future experiments will be required to determine if buprenorphine activation of the NOP receptor during development can affect adult NOP receptor function and lead to the development of psychiatric disorders.

The current results suggesting that buprenorphine can alter cortical development are in line with human studies that demonstrate that buprenorphine exposure *in utero* may have long-term developmental consequences. Specifically, 5-6 year old children exposed to buprenorphine *in utero* have been shown to have deficits in motor skills and memory. These children also had increased hyperactivity, impulsivity, and attention problems compared to peers matched for neonatal abstinence syndrome, gender, and socio-economic factors (15). Other studies, however, have failed to demonstrate that prenatal buprenorphine exposure produces deficits in cognitive development, language abilities, sensory processing, and temperament over the first 3 years of life (52). These contradictory findings in humans may be a result of the numerous potential confounding factors at play, including the severity of neonatal abstinence syndrome and obstetric complications (e.g. low birth weight, etc.) as well as the home environment. Further, these studies are limited to infants and young children. However, the current studies suggest that buprenorphine exposure *in utero* may influence circuits involved in psychiatric disorders, which often do not present until adulthood. There are currently no human studies that follow buprenorphine exposed babies into adulthood. However, rodent studies show that prenatal buprenorphine exposure can decrease neurogenesis and increases depression- and anxiety-like behavior in adult animals (53, 54), suggesting that buprenorphine may have long-term consequences.

In conclusion, a large body of research indicates that OUD can have severe and immediate consequences on a fetus, including low birth weight, sudden infant death, respiratory complications and Neonatal Abstinence Syndrome (14). Buprenorphine has been shown to improve outcomes in infants, decreasing the incidence, severity, and duration of Neonatal Abstinence Syndrome compared to mothers using illicit opioids (7) and those that receive other treatments for OUD (55, 56). Therefore, our results should not be interpreted as a reason to stop the use of buprenorphine in pregnant women with OUD. However, our findings suggest that more research should be done to increase our understanding of the long-term consequences of buprenorphine exposure *in utero*, and to find novel therapeutics, particularly those that do not target the NOP receptor.

## Acknowledgements

This work was supported by a SALSI pilot award for opioid epidemic research (to J.H. and D.L.), NIH grants (R01NS093992, R01NS113516, R01NS089770, and R21AG066496 to J.H., MH090067 to D.L., K99MH121355 to J.D.), the VA (BX004693 and BX004646 to D.L.), and the Robert J. Kleberg, Jr. and Helen C. Kleberg Foundation and the Semmes Foundation (to J.H.). We thank Aline McKenzie for manuscript editing. Some figures were created with BioRender.com.

---

**Supplementary Figure 1.**
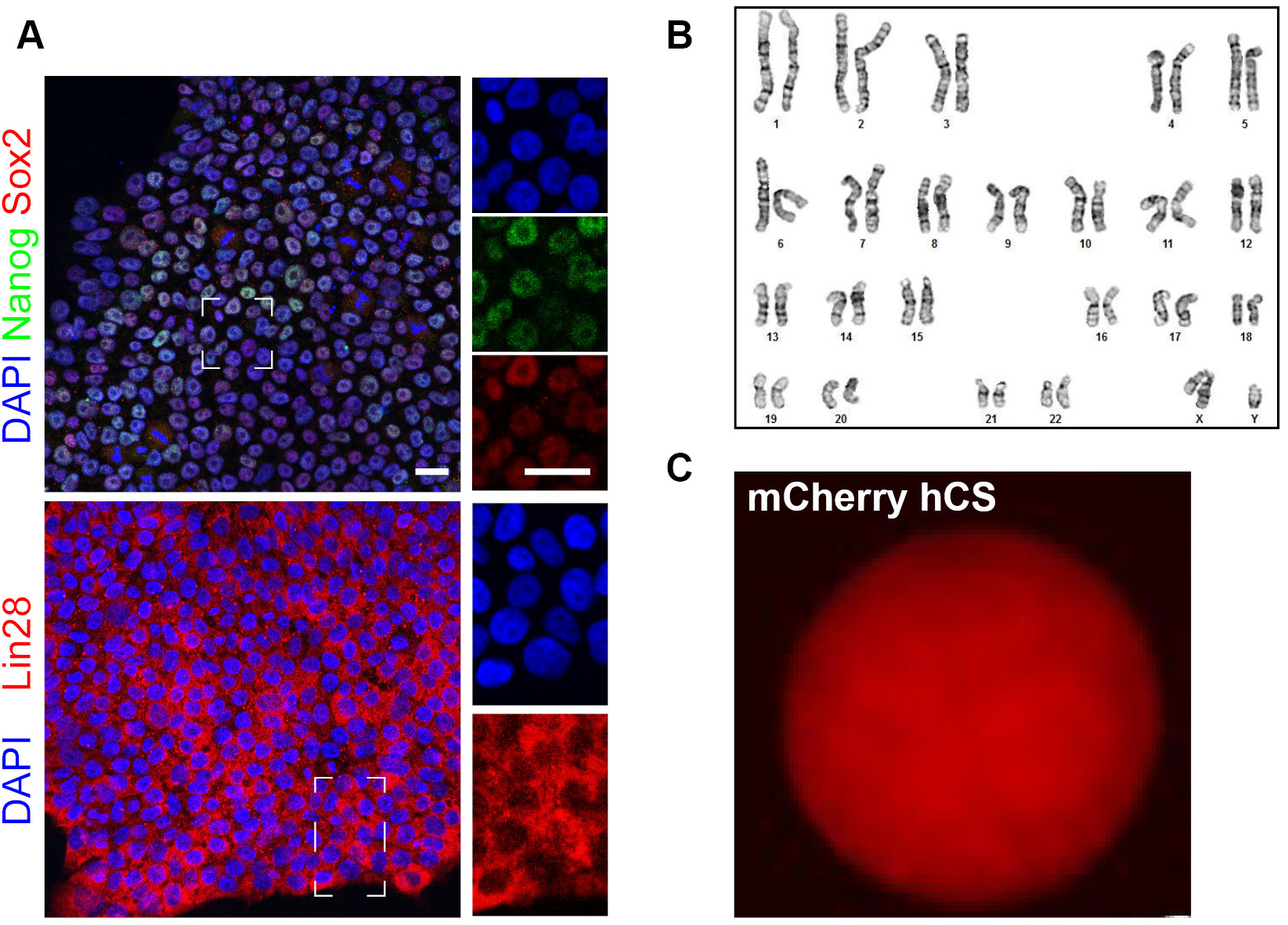
A) hiPSC lines maintained markers of pluripotency. B) hiPSC lines maintained a normal karyotype. C) A representative image of a cortical spheroid generated from the CRISPR-mCh hiPSC cell line. Scale bar = 25 um

**Supplementary Figure 2.**
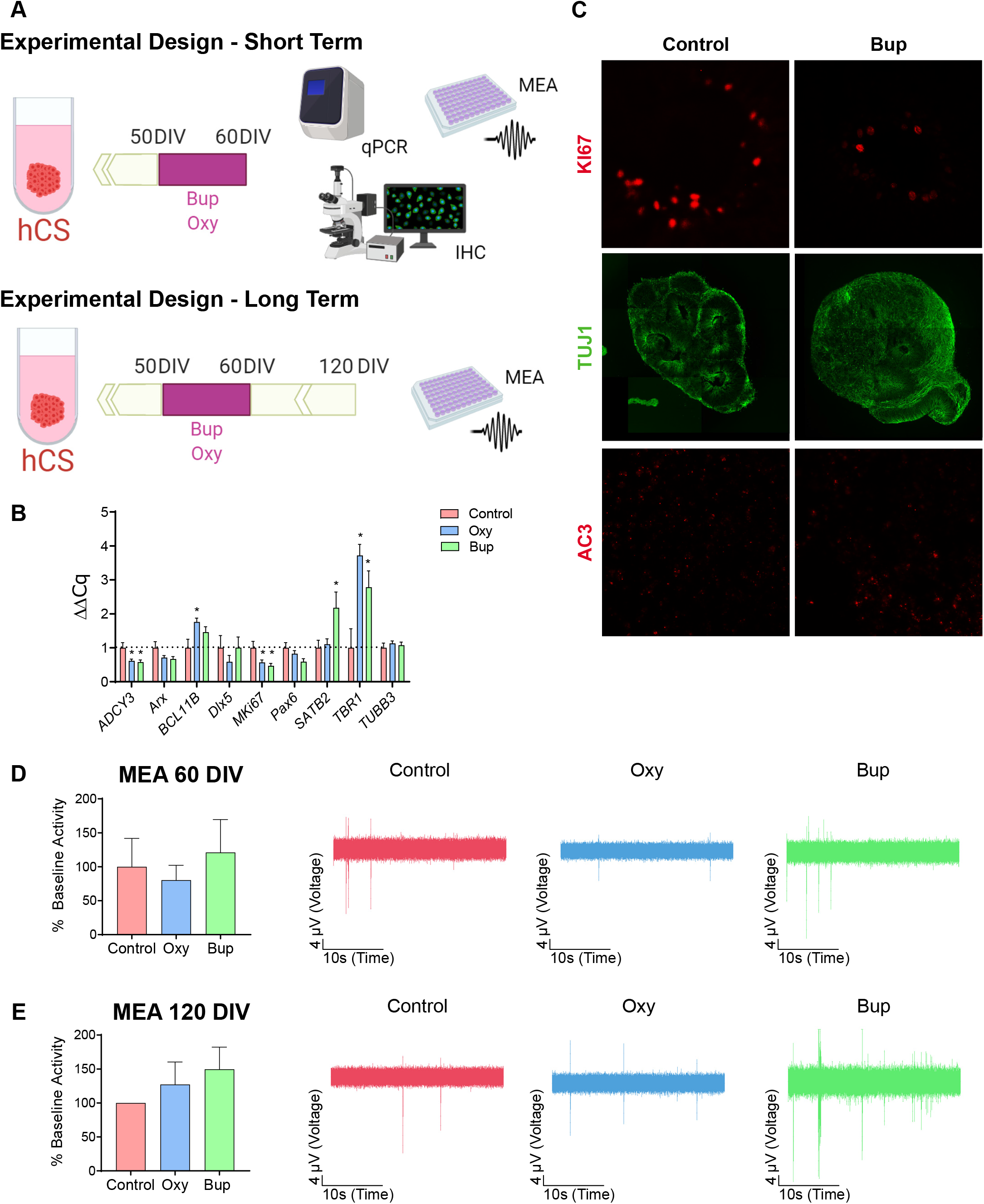
The experimental design is depicted in (A). B) Buprenorphine and oxycodone had some effects on markers of cortical development, as measured by qPCR. n=8. C) Representative images of Ki67, TUJ1, and AC3 immunohistochemistry in control- or buprenorphine-treated spheroids. D) Neither oxycodone or buprenorphine affect cortical network activity in MEAs performed on day 60. n=20-24. E) Neither oxycodone or buprenorphine affect cortical network activity in MEAs performed on day 120. n=25. Scale bar = TBD

